# Human microbiome sequences in the light of the Nubeam

**DOI:** 10.1101/763631

**Authors:** Hang Dai, Yongtao Guan

## Abstract

We present Nubeam (nucleotide be a matrix) as a novel reference-free approach to analyze short sequencing reads. Nubeam represents nucleotides by matrices, transforms a read into a product of matrices, and based on which assigns numbers to reads. Nubeam capitalizes on the non-commutative property of matrix multiplication, such that different reads are assigned different numbers, and similar reads similar numbers. A sample, which is a collection of reads, becomes a collection of numbers that form an empirical distribution. We demonstrate that the genetic difference between samples can be quantified by the distance between empirical distributions. Nubeam can account for GC bias and nucleotide quality, and is computationally efficient; the K-mer method is a special case of Nubeam, but without those benefits. As a reference-free approach, Nubeam avoids reference bias and mapping bias and can work with organisms without reference genomes. Thus, Nubeam is ideal to analyze datasets from metagenomic whole-genome sequencing, where the amount of unmapped reads is substantial. When applied to human microbiome sequencing, Nubeam recapitulated findings made by mapping-based methods, and shed lights on contributions of unmapped reads. In particular, body habitats dominate clustering of unmapped pseudo-samples; there are more outliers in skin whole samples than the skin mapped pseudo-samples; and analysis of unmapped reads suggested that the sequencing depth is far from sufficient for urogenital samples.

## Introduction

When identifying variants is not a must and the primary interest is to quantify genetic differences between samples (Ravel et al., 2011; Nayfach and Pollard, 2016), it can be beneficial to analyze short sequencing reads without reference genomes. First, it avoids reference bias and mapping bias. Both biases can be alleviated but never overcome because they are intrinsic to the mapping based approach. Second, it avoids uncertainty related to variants-call, particularly when the sequencing coverage is low. Third, it becomes possible to analyze organisms that have no reference genomes, or the reference genomes are incomplete or in low quality.

The prominent reference-free approach is the K-mer method (Jiang et al., 2012; Subramanian and Schwartz, 2015; Lu et al., 2017). Simply put, the K-mer method calculates frequencies of each K-mer (K consecutive nucleotides) presented in all reads from a sample, and infer differences between samples by comparing K-mer frequencies. In practice, however, the K-mer method has several difficulties. First, it implicitly assumes error-free in reads, and it is difficult—if not impossible—to account for nucleotide quality (Comin et al., 2015; Comin and Schimd, 2016). Second, choosing K can be a headache—too small or too large of K will make the K-mer frequencies less informative. Third, some pairs of K-mers only differ by one nucleotide and other pairs of K-mers differ by K nucleotides, but it is difficult to account for such differences in the K-mer method. Last but not least, how to account for GC bias is an unmet challenge for the K-mer method.

We present a novel method, Nubeam, that includes the K-mer method as a special case, but with several key advantages. Nubeam can account for nucleotide quality and GC bias, and its computation is efficient. As a reference-free method, Nubeam is particularly suitable to analyze sequencing data in metagenomics, because it is a norm that a large percent (on average more than half) of the reads cannot be mapped to reference genomes (Supplementary Figure S1). We apply Nubeam to analyze sequencing reads from the human microbiome project (Consortium et al., 2012) and our main objective is to understand how the unmapped reads affect the genetic distance estimates between samples.

## Results

### Nucleotide be a matrix — Nubeam

We assumed each sample is a collection of reads of the same length *l*, and each read consists of four nucleotides A, T, C and G, and possibly an unknown nucleotide denoted by N. We first constructed four binary sequences from a read by using each of the four nucleotides as a reference, with the reference nucleotide being 1 and the others 0. (Note that our representation allows us to mask a low quality nucleotide to N to account for nucleotide quality.) We used a matrix 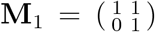 to represent 1, and its transpose 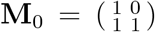 to represent 0. For each binary sequence *B* = *b*_1_*b*_2_ … *b_l_* we obtained the product matrix 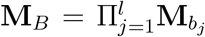. (When B is an alternating sequence 101010…, the entries in **M_*B*_** are entries in Fabonacci sequence.) Let **W** be a weight matrix, we designated log(*tr*(**WM_*B*_**)) as the Nubeam number to the binary sequence *B*. (Here we used 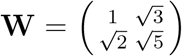) and this choice will become clear below.) Thus, for each read we obtained four Nubeam numbers (Nubeam quadruplet) that jointly represent the read (Figure 1a).

**Figure 1:**
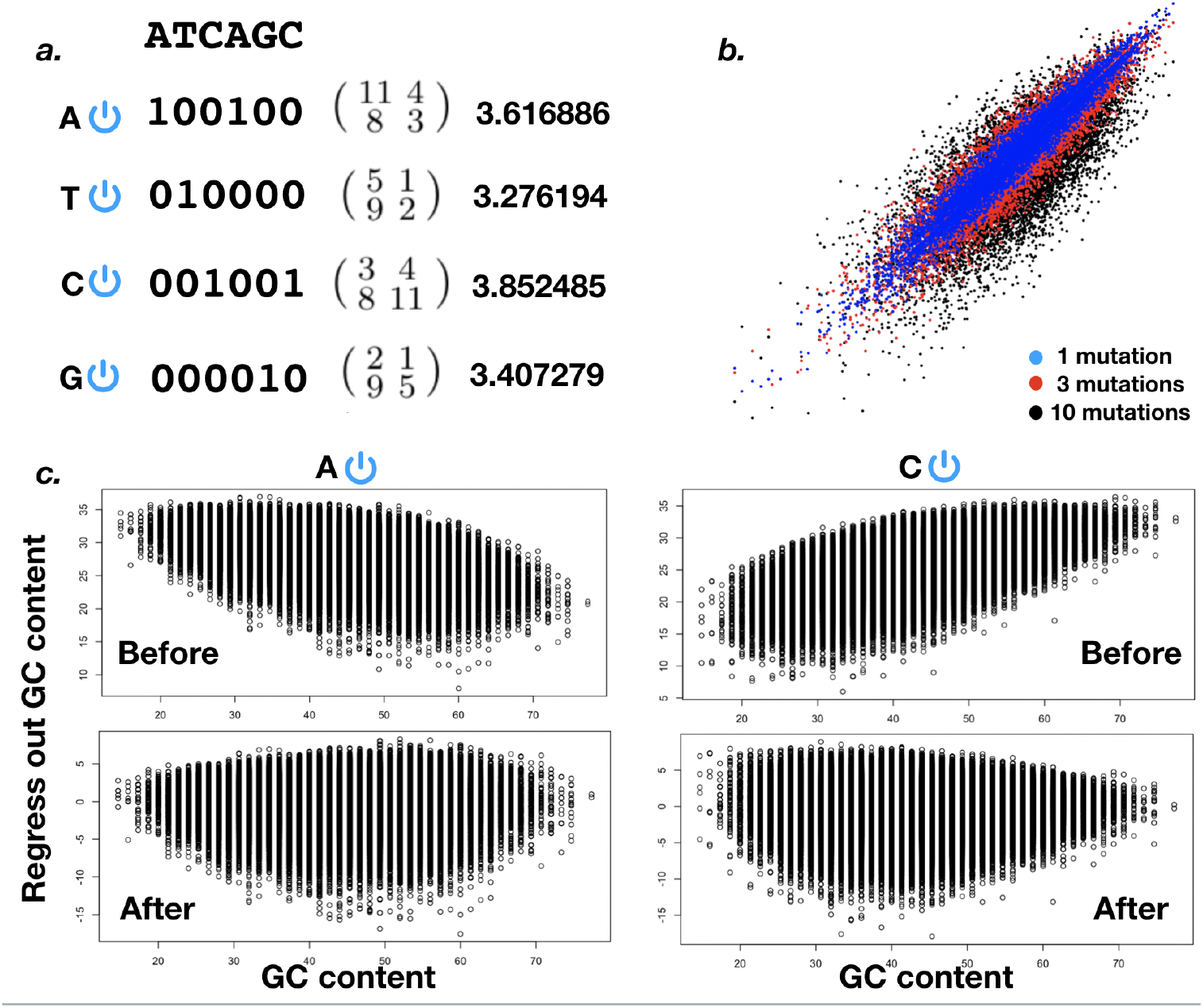
a. Cartoon of how to obtain Nubeam quadruplet for a read. Convert a read to four binary sequences, turn each binary sequence into a product matrix, and obtain a number from each product matrix. b. Similar binary sequences produce similar numbers. For each simulated binary sequence of length 100, we obtained sequences with 1 or 3 or 10 random mutations, and compare the Nubeam numbers between original sequences and their mutant sequences. c. Regress out GC content from Nubeam numbers of binary sequences. Left: with A as reference (T as reference is similar). Right: with C as reference (G as reference is similar).

For each sample, a collection of reads becomes a collection of Nubeam quadruplets. These quadruplets define a four dimensional empirical distribution, and the genetic distance between two samples can be quantified by the distance between two empirical distributions. We obtain histogram estimates for each collection of quadruplets (Supplementary Material and Methods), and calculate the Hellinger distance between probability mass functions (Methods). We will show that the Nubeam Hellinger distance is highly correlated with genetic distance between two samples.

Nubeam capitalizes on the non-commutative property of matrix multiplication, so that if two reads differ, their Nubeam quadruplets differ. More importantly, if two reads are similar (e.g. small Hamming distance), their Nubeam quadruplets are close to each other. To demonstrate this, we simulated binary sequences of length 100, mutated (by flipping the digit) at 1, or 3, or 10 random positions to obtain mutant sequences, and compared their Nubeam numbers. Figure 1b plotted Nubeam numbers of original binary sequences (x-axis) vs mutant sequences (y-axis) with different number of mutations. Indeed, similar sequences tend to have similar Nubeam numbers. This “continuum” property is important as it is the foundation for the usefulness of the Nubeam representation. The effectiveness of quantitative treatments, including controlling for GC bias and histogram estimates for empirical distribution, implicitly depends on the continuum property.

The usefulness of a method depends on how effectively it handles the known artifacts in real data. One major artifact in sequencing is strand bias, where one strand may be sequenced in a much higher proportion than the other strand. Here we provide a simple solution to account for strand bias. For a read R we obtain its reverse complement read U, and we compute quadruplets for both R and U for building an empirical distribution. The other major artifact in sequencing is the GC content bias, mainly due to the intrinsic property of the polymerase chain reaction (PCR), leading to the fact that some stretch of nucleotides are more likely to be sequenced than others depending on the GC content (Benjamini and Speed, 2012). We control GC bias by regressing out GC counts from each quadruplets using the simple linear regression, treating each read as unit (Figure 1c). We found it more effective to perform regression jointly over all samples, instead of one sample at a time. After GC being regressed out, the residuals are used to define empirical distribution to quantify genetic distances between samples.

### Nubeam distance reflects genetic distance

We first demonstrate that Nubeam distances correlate well with the genetic distances in a three-taxa setting (Figure 2). Let a star tree have three leaf nodes and the inner node is seeded with a substrain of *Escherichia. coli.* Taxa S1 and S2 have equal branch lengths that correspond to 1 × 10^-5^ mutation per site per cell division. We varied the branch length of taxa S3, with the corresponding mutation rate ranging from 1 × 10^-5^ to 1 × 10^-3^ per site per cell division. In perspective, two phenotypically similar *E. coli* strains from different environments differ by about 1000 nucleotides (Swick et al., 2013), which correspond to mutation rate of 1 × 10^-4^. This simulation setup produces varying genetic distances between S3 and S1/S2 with fixed distance between S1 and S2 as a baseline. We then simulated genomes of S1, S2, and S3 using iSG (Strope et al., 2006) under the general time reversible (GTR) substitution model (Tavare, 1986; Setti et al., 2012). For each simulated genomes, we produced 75bp error-free reads using a sliding window. The simulation were replicated 100 times. Figure 2 (a) plotted the true genetic distance (measured by the Hamming distance normalized by the genome size) vs inferred Nubeam distance, and the Spearman’s rank correlation between the Nubeam distance and genetic distance is 0.999.

**Figure 2:**
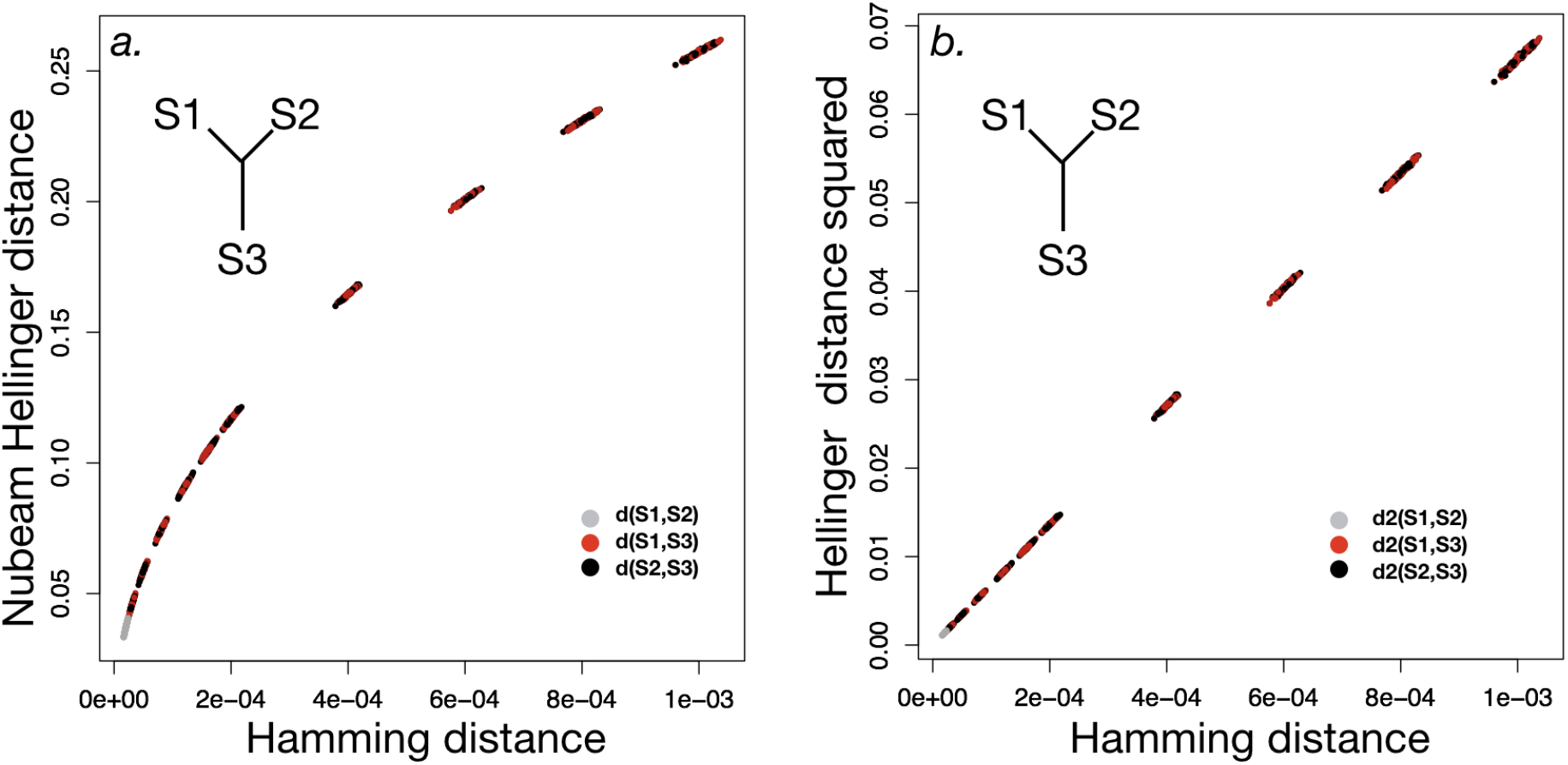
Nubeam Hellinger distance correlates well with genetic distance. a. Hamming distance (x-axis, normalized by genome size) v Nubeam Hellinger distance (y-axis). b.The squared Hellinger distance (y-axis) appears to be linear with the Hamming distance (x-axis).

Next we demonstrated that true phylogenies could be reconstructed using Nubeam distance. We simulated sets of sequencing reads using eight-taxonomic-unit trees (Supplementary Material and Methods), computed Nubeam distances between each pair of taxa to reconstruct the phylogenetic tree, and assessed the accuracy of phylogeny reconstruction by comparing the inferred tree with the true tree (Chan et al., 2014). We used six combinations of internal and terminal branch lengths of 1 × 10^-5^ and 5 × 10^-5^ to represent different degrees of genomic difference and different levels of difficulties for phylogeny inference (Supplementary Figure S2). For each of the 100 replicates of a tree, hierarchical clustering was applied on the Nubeam distance matrix to reconstruct the phylogeny. The reconstructed phylogenies were compared with the true tree using Compare2Trees (Nye et al., 2005), and Nubeam perfectly reconstructed phylogenies for each of 100 replicates for all six generating trees.

### Nubeam controls for GC bias

Based on the *E. coli* genome, we simulated 15 samples without GC content bias and 30 samples with GC content bias, 10 for each bias scheme. The bias schemes are detailed in the Methods section, and the severity of the GC bias in each schedule can be observed in Figure 3 (a). These three schemes reflected typical GC bias patterns we observed in whole genome sequencing data for humans. Let *d*_0_ represent the Nubeam distance between two samples without GC bias, let *d_b_* represent Nubeam distance between a sample without GC bias and a sample with GC bias, but without controlling for GC bias, and let *d_c_* represent Nubeam distance between a sample without GC bias and a sample with GC bias, but controlling for GC bias. To see the effectiveness of Nubeam controlling for GC bias, we compared *r_b_* = (*d_b_* – *d*_0_)/*d*_0_ and *r_c_* = (*d_c_* – *d*_0_)/*d*_0_, both visually and numerically (Figure 3 b,c,d). The GC biases were reduced dramatically and effectively in all three GC bias simulation schemes.

**Figure 3:**
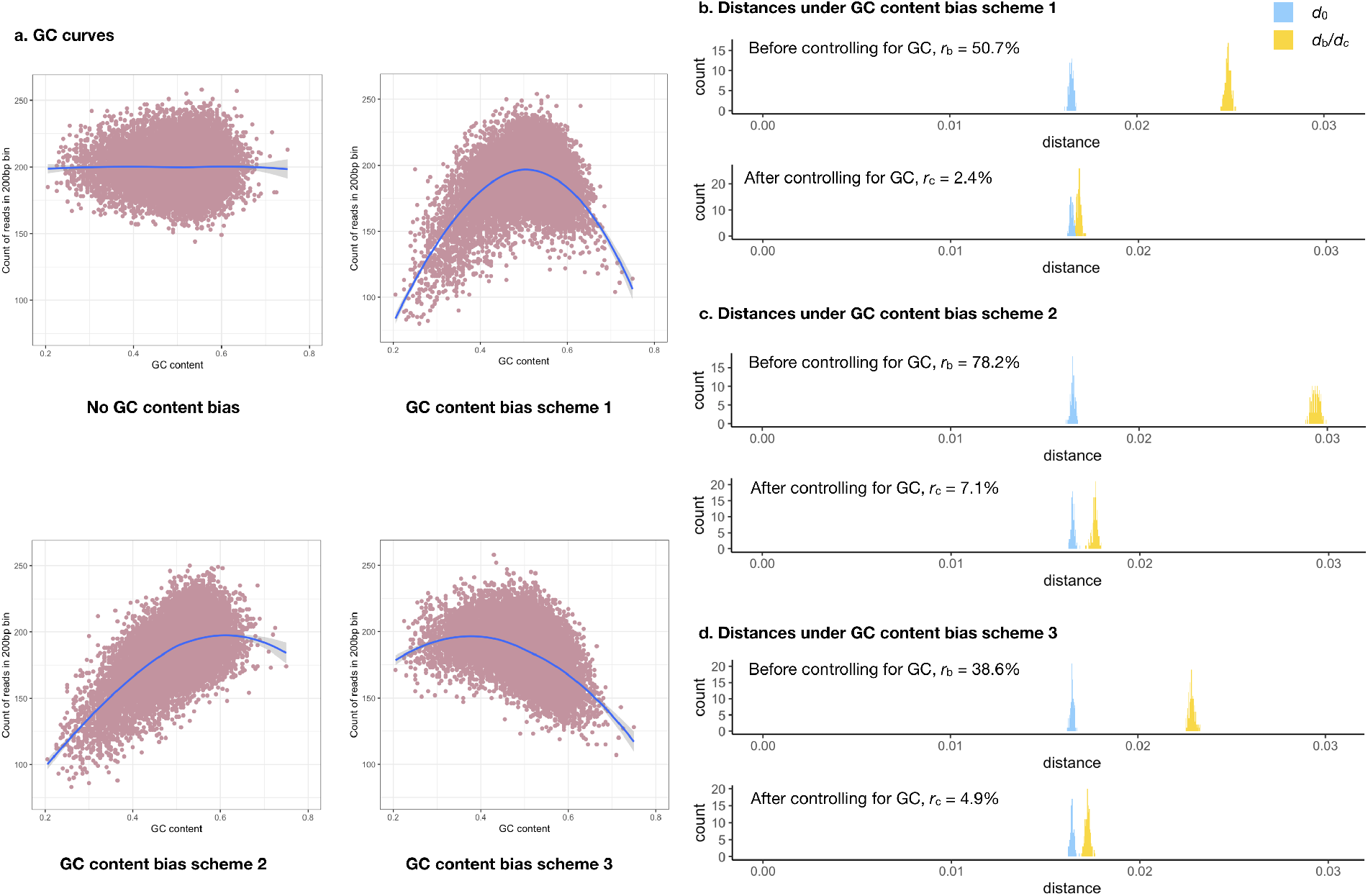
The effect of controlling for GC content bias in Nubeam. (a) GC (x-axis) vs bin coverage (y-axis) to demonstrate patterns of GC bias for three simulation schemes with LOESS curve marked in blue. Recall 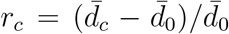 and 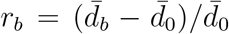 measure relative difference before controlling for GC bias and after controlling for GC bias. b) *r_c_* = 2.4% after controlling for GC bias and *r_b_* = 50.7% before controlling for GC bias for simulation scheme 1. (c) *r_c_* = 7.1% after controlling for GC bias vs *r_b_* = 78.2% before controlling for GC bias for simulation scheme 2. (d) *r_c_* = 4.9% after controlling for GC bias vs *r_b_* = 38.6% before controlling for GC bias for simulation scheme 3.

### Nubeam includes the K-mer method as a special case

We first show that if two binary sequences are different then their corresponding product matrices differ (Proposition 1). By definition of 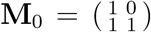 and 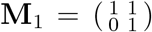, a matrix right multiplying an **M**_0_ is to add the second column of the matrix to the first, and right multiplying an **M**_1_ is to add the first column to the second. Thus for a given product matrix, one can solve for the binary string using the following algorithm. Let the binary string be *b*_1_*b*_2_ … *b_l_*. Compare the columns of the product matrix, pick the large column, if it is the first column then *b_l_* = 0 and if it is the second then *b_l_* = 1. Subtract the small column from the large column to obtain a new matrix, let *l* decrease by 1 and repeat the procedure.

Because the algorithm is deterministic, the solution must be unique. Thus, two different binary sequences must correspond to two different product matrices. Let **F** and **G** be two matrices whose entries are integers, and recall 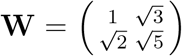, then *tr*(**WF**) = *tr*(**WG**) if and only if **F** = **G** (Proposition 2). This is true because any entry in 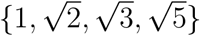 is not a linear combination of the other three entires with rational coefficients (Boreico, 2008).

If two reads are different, then at least one pair of binary strings (among four such pairs) are different, and consequently (by Proposition 2) these two binary strings get assigned different numbers. Thus, if two reads are different then the two quadruplets are different. When we use each unique quadruplet as a bin to compute distance between empirical distributions, we obtain the K-mer method. Compared to Nubeam, the K-mer method is rather primitive: it has difficulty to control for GC bias and to account for sequencing error, it disregards the different degree of similarities among K-mers, and its computation becomes difficult rather quickly as *K* increases.

### Nubeam analyzing human microbiome project data

Metagenomics data from whole genome shortgun-sequencing (WGS) is ideal for Nubeam, simply because there are more than half reads cannot be mapped anywhere in more than 3000 microbial reference genomes. We analyzed 690 samples from 7 body habitats (excluding a habitat called “other oral”) of phase I data from the human microbiome project (HMP) (Consortium et al., 2012). HMP performed quality control and removed human sequence contamination; reads passing low-complexity filter were aligned to reference genomes, with 57.28 ± 13.39% of reads per sample mapped (Consortium et al., 2012) (Supplementary Figure S1). The inadequacy of the reference genomes and mapping bias both contribute to the high percentage of unmapped reads (Nayfach and Pollard, 2016). We split each sample (whole-sample) into mapped reads (mapped pseudo-samples) and unmapped reads (unmapped pseudo-samples). Our analysis has two aims: whether we can recapitulate the findings from mapping based method using Nubeam to analyze mapped reads, and whether Nubeam enables the unmapped reads to provide additional information.

#### Nubeam within-sample diversity

We used the second moment of Nubeam numbers of a sample to quantify within-sample diversity (see Methods). To validate our definition of alpha-diversity, we simulated a mixture of two bacteria species (*E. coli* and *Proteus. mirabilis*) by mixing reads at various mixing proportions. The inferred within-sample diversities reflected our intuition such that the intermediate mixing proportions have higher alpha-diversity than extreme mixing proportions (Supplmentary Figure S4).

We quantified within-sample diversity of mapped and unmapped pseudo-samples using Nubeam. Being able to quantify within-sample diversity of the unmapped samples is a strength of Nubeam. For mapped pseudo-samples, oral and gastrointestinal habitats have high within-sample diversity, whereas skin and vagina have low within-sample diversity (Figure 4)—the ranking of within-sample diversity by body habitats is largely consistent with that in published study (Consortium et al., 2012) of HMP. For unmapped pseudosamples, the within-sample diversity is generally higher than mapped ones, particularly for nasal and urogenital samples (Figure 4); the ranking by body habitats is also different from mapped ones. Interestingly, there is strong linear relationship between within-sample diversity of mapped and unmapped pseudo-samples for most body habitats except buccal mucosa and vagina.

**Figure 4:**
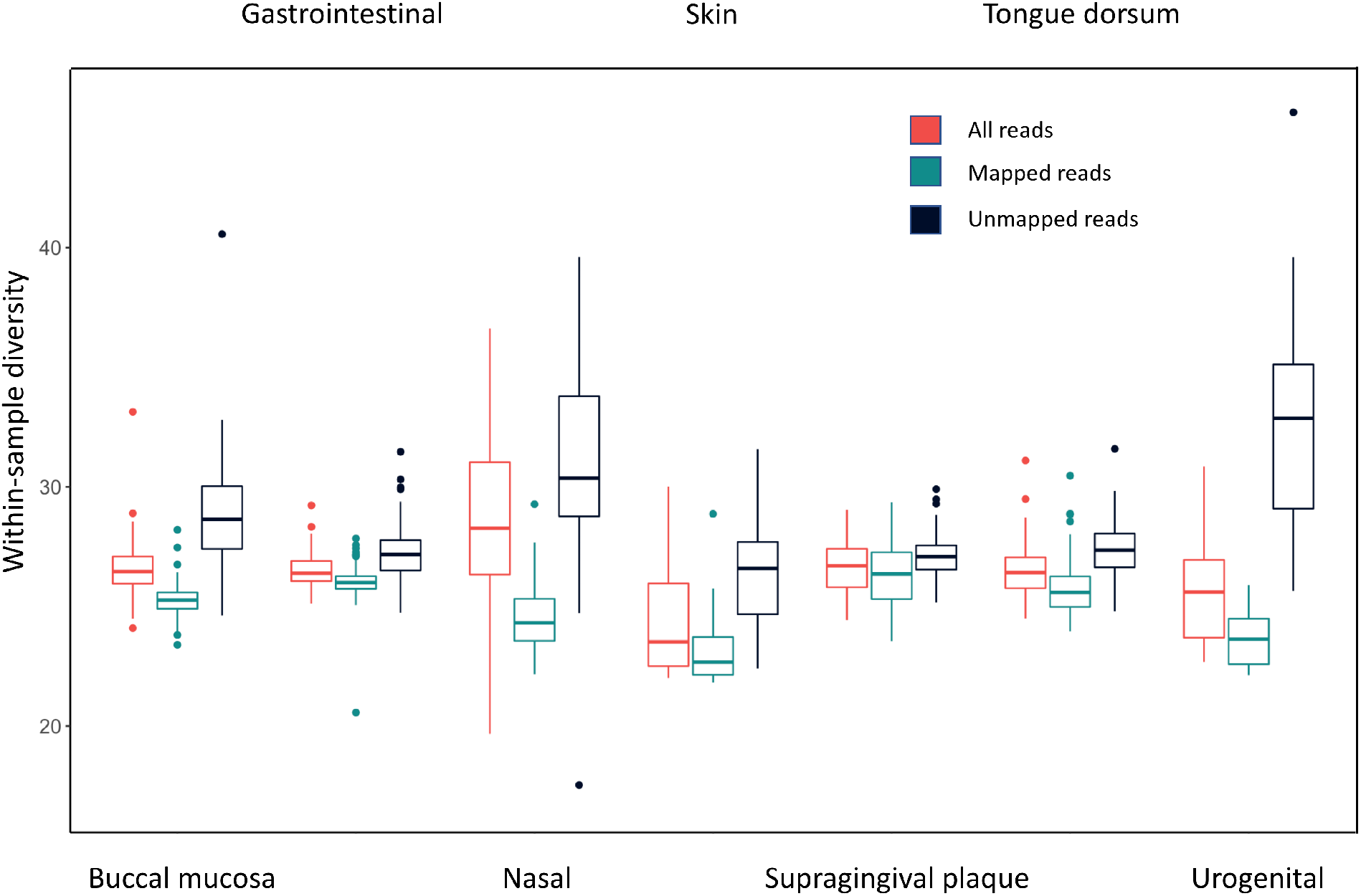
The within-sample diversity of the whole sample, and mapped, unmapped pseudosamples for each body habitats.

#### Nubeam between-sample distance

In metagenomics literature, the beta-diversity measures how different of microbial composition between two samples. Here we used Nubeam distance to quantify beta-diversity between a pair of samples, and examined beta-diversity estimates using hierarchical clustering. In parallel, we applied UMAP (Maaten and Hinton, 2008) for non-linear dimensionality reduction and visualization to examine beta-diversity estimates (Supplementary Figure S3). UMAP clustering corroborates hierarchical clustering on big pictures, but for finer details such as outliers we trust hierarchical clustering over UMAP.

We first examined the beta-diversity of mapped pseudo-samples (Figure 5 (a)). The primary clustering was by body habitats, with gastrointestinal, oral and urogenital samples well separated. Skin samples were intermingled with a subset of nasal samples. Within oral cavity, samples from supragingival plaque were clearly separated from those from buccal mucosa and tongue dorsum, and a proportion of buccal mucosa samples were intermingled with tongue dorsum samples. These results were comparable with published studies of the same datasets using mapping-based methods (Consortium et al., 2012).

**Figure 5:**
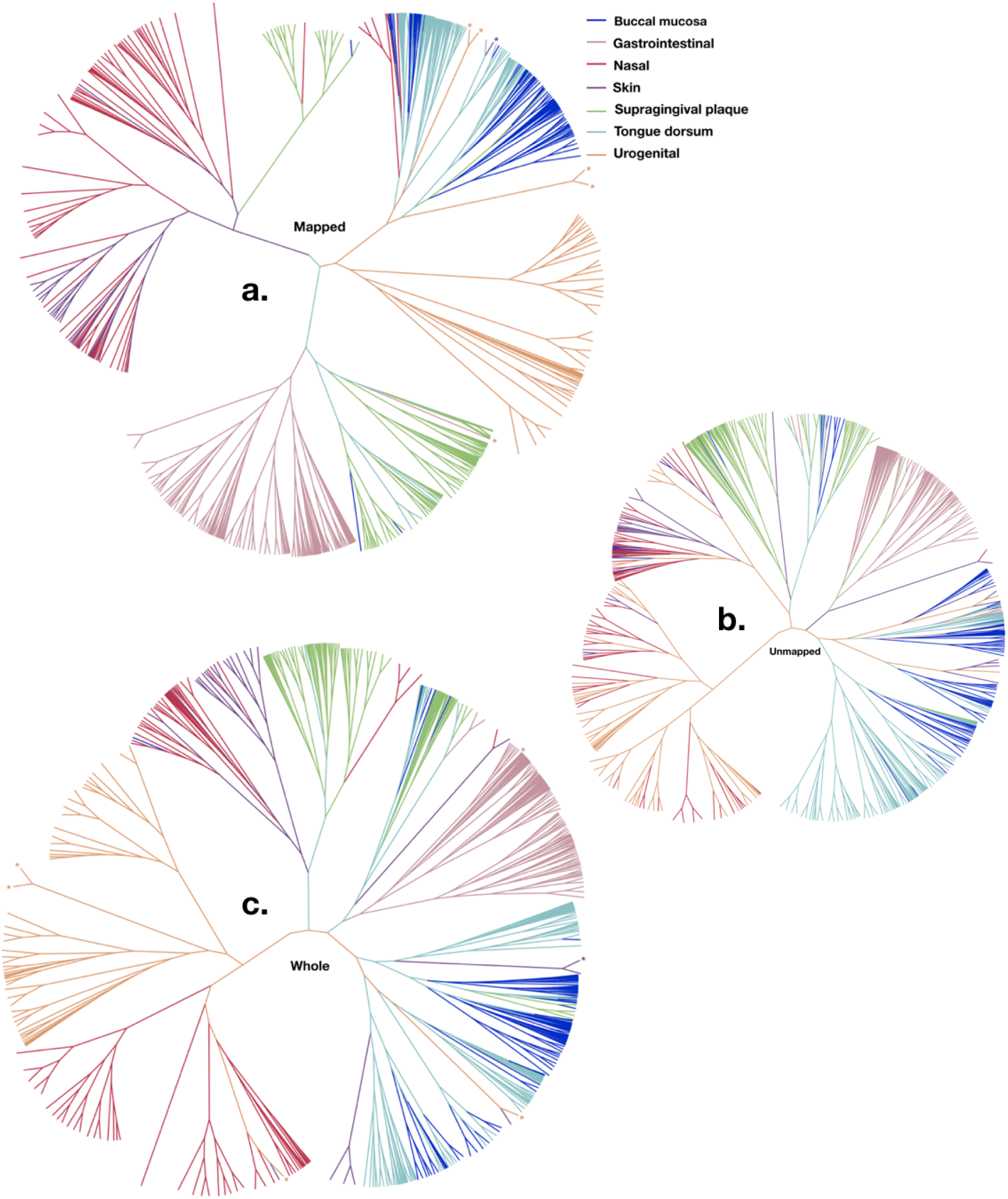
Samples clustering for mapped pseudo-samples, unmapped pseudo-samples, and whole samples respectively by hierarchical clustering. If a sample is marked by a * in (a), then its corresponding whole sample was again marked by a * in (c).

We then examined the beta-diversity of unmapped pseudo-samples (Figure 5 (b)). Re-assuringly, the overall clustering pattern still follows the body habitats. The most striking difference is that vaginal samples intermingled with the nasal samples, which is also evident in Supplementary Figure S3 (b). Lastly, we examined the beta-diversity of whole samples. Figure 5 (c) showed that the clustering of whole samples are also by body habitats, largely agreed with those of mapped pseudo-samples. Noticeable differences do exist, however, presumably due to unmapped reads. For example, there were only 2 outliers for skin in mapped pseudo-samples, but 8 outliers for skin in whole samples; one Gastrointestinal outlier for mapped pseudo-samples (marked by * in Figure 5 (a)), distinguished by its unusually high contents (7.5%) of pathogen *Shigella* spp., was no longer a outlier in whole samples; one skin outlier was clustered with oral mapped pseudo-samples, likely due to its 46.1% of *Finegoldia magna,* an opportunistic pathogen that can be found in skin, oral, gastrointestinal and urogenital habitats (Rosenthal et al., 2012), and its corresponding whole sample was still an outlier but with a new companion.

#### Urogenital (vaginal) samples

There were four urogenital outliers in the mapped pseudo-samples (Figure 5 (a)). To make sense these four samples, we compared the clustering with the taxonomic compositions of all urogenital samples (Methods). Figure 6 clearly demonstrated that the clustering is in concordance with their taxonomic compositions and abundances. In particular, two outlier samples have high proportions of anaerobic bacteria dominated by *Gardnerella* and *Atopobium,* but no *Lactobacillus* bacteria; the other two outlier samples contain a large proportion of *Bifidobacterium* bacteria (10% and 31% for the two samples respectively), in addition to a high proportion of *Lactobacillus gasseri*.

**Figure 6:**
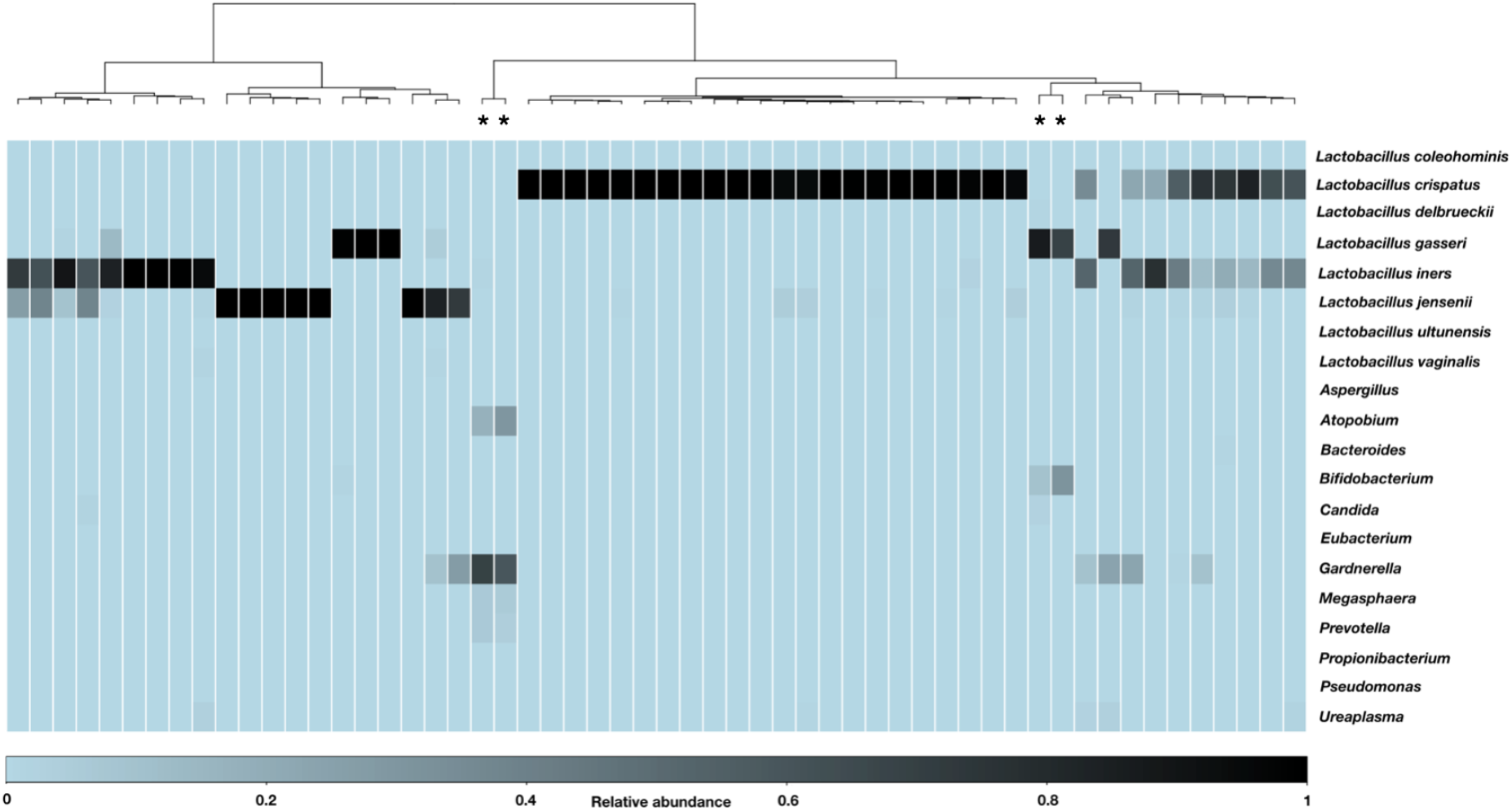
Hierarchical clustering of vaginal samples using Nubeam calculated distance matrix is consistent with community state types defined by relative abundance of microbial taxa. The heatmap was generated using relative abundances of microbial taxa, with only the most abundant ones were chosen. The four outlier samples are marked by star. Ward’s minimum variance method was used for hierarchical clustering.

Vaginal microbiomes are known to have simple taxonomic compositions dominated either by a single *Lactobacillus* species or by strictly anaerobic bacteria (Ravel et al., 2011; Romero et al., 2014). But our analysis suggest that might be only true for mapped pseudosamples, as the unmapped urogenital pseudo-samples has extraordinarily high within-sample diversities. We thus performed de novo assembly using MEGAHIT (Li et al., 2015) for all unmapped reads to produce contigs (minimum length 228 bp) and remapped reads to contigs. To our surprise, 74% of the reads failed to be re-mapped, indicating they are isolated reads; 18% reads can be mapped to contigs with null BLAST results, indicating they are from unknown organisms; only 8% reads can be mapped to contigs from known micro-organisms (according to BLAST results). These results suggested that the sequencing depth for vaginal samples in HMP is far from sufficient, and the composition of vaginal microbiome need to be further studied using WGS with sufficient coverage.

## Discussion

We presented a novel reference-free approach Nubeam to analyze short sequencing reads. Nubeam can account for sequencing error, strand bias, and GC bias to quantify genetic distance between samples. We showed that the K-mer method is a special case of Nubeam, stripping off its ability to account for sequencing error and GC bias. We demonstrated its usefulness by applying Nubeam to analyze datasets from whole-genome sequencing in HMP. Our analysis shed new lights on the HMP in several aspects. First, similar to the clustering of the mapped pseudo-samples, the clustering of unmapped pseudo-samples is aslo dominated by the body habitats. Second, the clustering differences between mapped pseudo-samples and whole samples do exist, and they are best represented by outlier samples of particular body habitats. Third, analysis of the unmapped reads suggested that the sequencing depth for vaginal samples is far from sufficient, and deeper sequencing might challenges the past believe that vaginal microbiomes have simple taxonomic compositions.

The unmapped reads are more prone to artifacts produced in DNA handling and sequencing. Since Nubeam is a reference-free approach, we paid extraordinary attention to guard against possible artifacts. We performed additional quality control on sequencing reads, and removed duplicates and reads that can be mapped to human genomes. The clustering of the unmapped pseudo-samples, which largely agreed with that of the mapped pseudo-samples, reassured us that our additional quality control procedure was effective, and our novel insights based on unmapped reads were not driven by data artifacts.

Many aspects of Nubeam invites continuing investigations. One aspect is to design better matrices to represent nucleotides for specific applications. For example, we discovered that matrix format of quaternion performs better in certain simulations. The other aspect is the computation. For example, computing Nubeam Hellinger distance between two (four dimensional) empirical distributions may be made more effective via shrinkage density estimates (Ma, 2017). We believe Nubeam can find a broad range of applications. One application we are currently developing is to perform reads deduplexing before mapping. Since a unique read is assigned a unique quadruplet, the deduplexing can be done efficiently. Nubeam can be effective in some areas where the K-mer approach is useful, for example, characterization of protein binding sequence motif (Newburger and Bulyk, 2009), characterizing CpG island by the flanking regions (Chae et al., 2013), and characterizing sequence feature for haplotype grouping (Navarro-Gomez et al., 2015).

A recent study suggested that environment, not genetics, primarily shapes the microbiome composition (Rothschild et al., 2018). Through quantifying microbiome difference between samples, Nubeam provides an effective method to quantify the environment factors to facilitates the study of gene environment interactions for many phenotypes such as diabetes and obesity. Finally, Nubeam enables us to study the contribution of unmapped reads to the genetic distance between samples. Using whole genome sequencing for human subjects, Nubeam has the potential to investigate whether and to what extent the unmapped reads can contribute to explain the “missing heritabilities” (Maher, 2008).

## Methods

### Controlling for GC bias

The GC content bias is a major sequencing artifact that leads to the dependence between regional coverage and GC content. When the signal of interest is the abundance of reads originating from certain genomic regions, GC content bias is a confounding factor. We correct the GC content bias at the read level by regression. We fit the standard linear regression model **y_i_** = **X***β* + *ϵ*, where **y_i_** is an *n* × 1 matrix of numbers assigned to reads by a matrix representation system. **X** is an *n* × 3 matrix of AT-count and GC-count of reads including a column vector of 1. *β* is a 3 × 1 vector of corresponding regression coefficients including the intercept; *ϵ* is an *n* × 1 vector of residuals. The residuals were then assigned to reads.

### Simulating GC content bias

We simulated sequencing samples with GC content bias using the following method (Benjamini and Speed, 2012). For a position *x* on the genome, the number of the 75bp read originating from *x* to be sampled follows Pois(λ), where λ is the expected count for the read. Denote the GC content (ranging from 0 to 1) of the 200bp fragment originating from *x* as *gc*. When there is no GC content bias, λ = 1 regardless of the value of *gc*; when there is GC content bias, λ is determined by Gaussian function 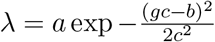. We simulated three schemes of GC content bias: *a* = 1, *b* = 0.5, *c* = 0.2 for scheme 1; *a* = 1, *b* = 0.6, *c* = 0.3 for scheme 2; *a* = 1, *b* = 0.4, *c* = 0.3 for scheme 3. These three schemes reflected typical GC bias patterns we observed in whole genome sequencing data for humans.

### Quantifying within-sample diversity

Let **X** be a collection of quadruplets. That is, **X** is an *n* by 4 matrix. Define **Σ** = **X**^*t*^**X**, and perform eigen-decomposition for **Σ**. The within-sample diversity can be quantified as the sum of eigenvalues (each eigenvalue is non-negative).

### Distance between two empirical distributions

Let X and Y be two collection of quadruplets. We divided samples into bins and obtained {*x_j_*} and {*y_j_*} as probability mass function for X and Y respectively. The Hellinger distance is defined as 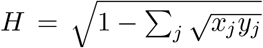. A detailed algorithm to partition bins in a balanced manner can be found in Supplementary Material and Methods.

### Human microbiome project samples

We downloaded bam files of 754 samples from http://hmpdacc.org/HMSCP/ and extracted reads using samtools. For a sample to be included in the study, it had to pass QC procedure described in http://hmpdacc.org/hmp/HMASM/, which leaves us 690 samples. Five samples had corrupted files and were removed. We further removed 29 samples from body site “other oral” and analyzed a total of 656 samples from 7 body sites. We deduplicated reads by an in-house software based on reads Nubeam numbers. Deduplicated reads were then mapped to human reference genome HG19 and human sequences from NCBI Nucleotide Collection (downloaded on Jun 7 2019) by BWA (Li and Durbin, 2009) to remove human contamination.

### Clustering of vaginal samples

For the 56 urogenital (vaginal) samples, we applied hierarchical clustering (with Ward’s minimum variance method) using Nubeam distance matrix. The heat-map of relative abundance of microbial taxa was generated using date by (Kraal et al., 2014), with the abundance tables of each sample downloaded from http://hmpdacc.org/HMSCP/. The abundance of strains in each sample was estimated by the product of breadth and depth of coverage, and its relative abundance was obtained by normalization. The relative abundance of species was calculated by adding up the relative abundance of strains belonging to same species, while the relative abundance of genus was calculated similarly. The selected strains/genera each had a cumulative relative abundance of more than 0.001 across 56 samples; and for each sample, their abundances accounted for more than 99.9% of overall abundance. *Lactobacillus* had all the species listed, while other genera only had their genus names listed.

## Data availability

The HMP data was downloaded from http://hmpdacc.org/HMSCP/, and the abundance tables of each sample was downloaded from http://hmpdacc.org/HMSCP/.

## Code availability

Nubeam source code is available on GitHub https://github.com/daihang16/Nubeam.

