## Supplementary Material and Methods, Supplemental Figures for "Human microbiome sequences in the light of the Nubeam"

### 1 Supplementary Material and Methods

#### 1.1 Simulation of genomes and inferences

We simulated genome for each taxonomic unit in the designed trees, with the root genome of *E. coli* strain K-12 substrain MG1655 (4,641,652 bp). We used iSG (Strope et al., 2006) for simulation under the general time-reversible (GTR) substitution model. The 6 parameters for relative substitution rate are as those estimated from the evolution study of bacteria rrsA, and 4 parameters for nucleotide frequency are those estimated from the E. coli genome. The branch lengths represent number of substitutions per basepair per generation. For the three-taxonomic-unit trees, both branch lengths of S1 and S2 had fixed values of $1 \times 10^{-4}$, while the branch length of S3 had values span two orders of magnitude from $10^{-2}$ to $10^{-4}$. For the eight-taxonomic-unit trees, we used six combinations of internal and terminal branch lengths of $1 \times 10^{-5}$ and $5 \times 10^{-5}$ to represent different difficulty levels of phylogeny inference.

Each simulation was done 100 replicates. For three-taxnomic-tree we only interested in relative distance between taxa. For eight-taxonomic-unit tree, we used hierarchical clustering (with Wards minimum variance method) to reconstruct trees from distance matrices produced by both Nubeam and the K-mer method. We compared the reconstructed trees with true trees by Compare2Trees (Nye et al., 2005), generating a score ranging from 0 to

*This work was partially supported by National Institutes of Health under award number R01HG008157. Most of work was done at Children's Nutrition Research Center of Baylor College of Medicine, the authors prior institution where HD was a PhD student and YG was an Assistant Professor. The work was also partially supported by United States Department of Agriculture/Agriculture Research Service under contract number 6250-51000-057.

1 for each comparison, with 0 indicating no topological similarity and 1 indicating same topology.

#### 1.2 Bin partitioning

The following algorithm does the balanced partition of bins.

**Algorithm 1** Balanced partition of bins using conditioning quantiles

```
Combine the matrices for all samples
Partition 1st column into BINs using quantiles
for each BIN of 1st dimension do
    for each ROW do
        collect the ROW if its 1st NUMBER is in BIN
    end for
    for the collected rows, partition their 2nd column into BINs usingquantiles
    for each BIN of 2nd dimension do
        for each ROW of the collected rows do
            collect the ROW if its 2nd NUMBER is in BIN
        end for
        for the collected rows, partition their 3rd column into BINs using quantiles
        ...
        for each BIN of (m − 1)th dimension do
            for each ROW of the collected rows do
                collect the ROW if its (m − 1)th NUMBER is in BIN
            end for
            for the collected rows, partition their m-th column into BINs using quantiles
        end for
    end for
end for
```

#### 2 Supplementary Figures

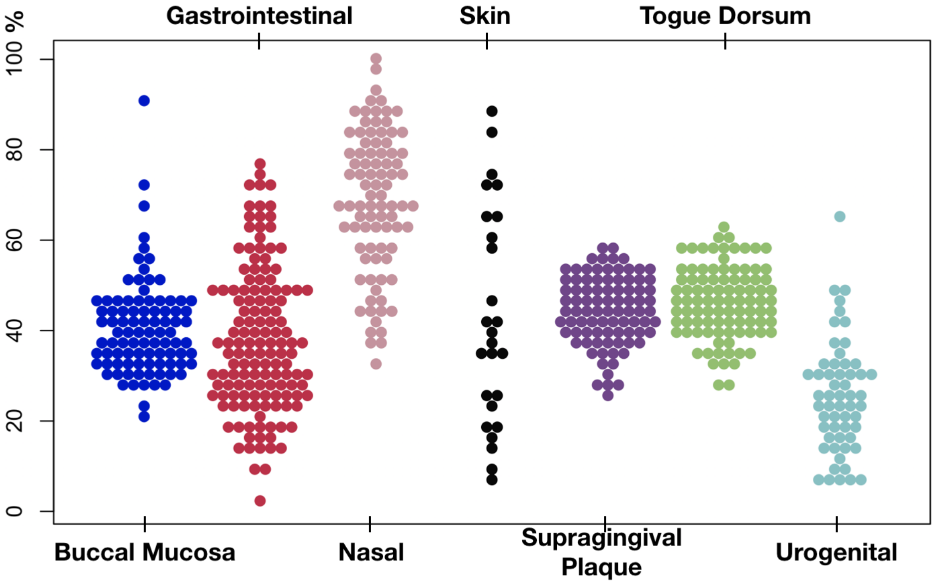


Figure S1: Percent of unmapped reads by body sites. Nasal samples have highest average unmapped reads, and gastrointestinal and urogenital samples have the lowest. Skin samples have large spread of unmapped reads.

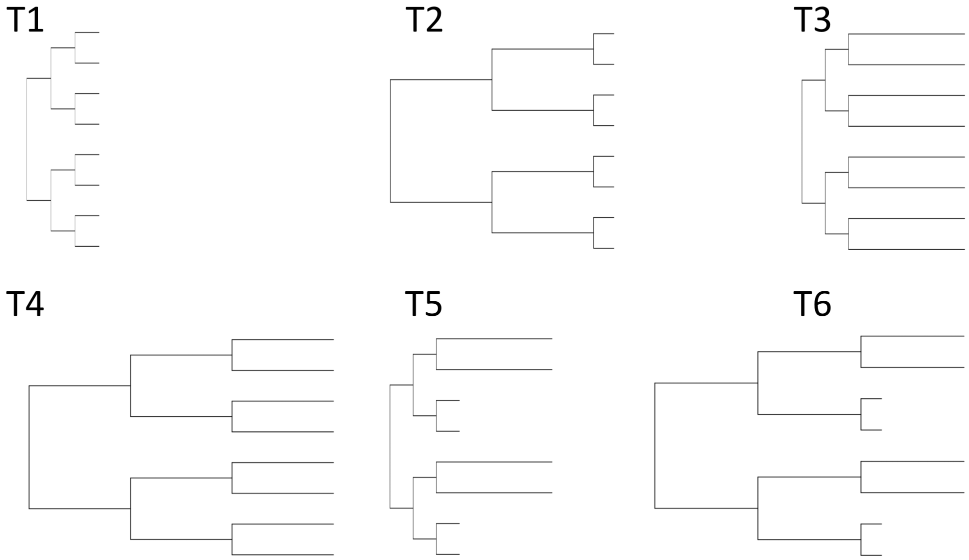


Figure S2: Six combinations of internal and terminal branch lengths of 0.0001 and 0.0005 were used to represent different degrees of genome divergence and thus different difficulty levels of phylogeny inference.

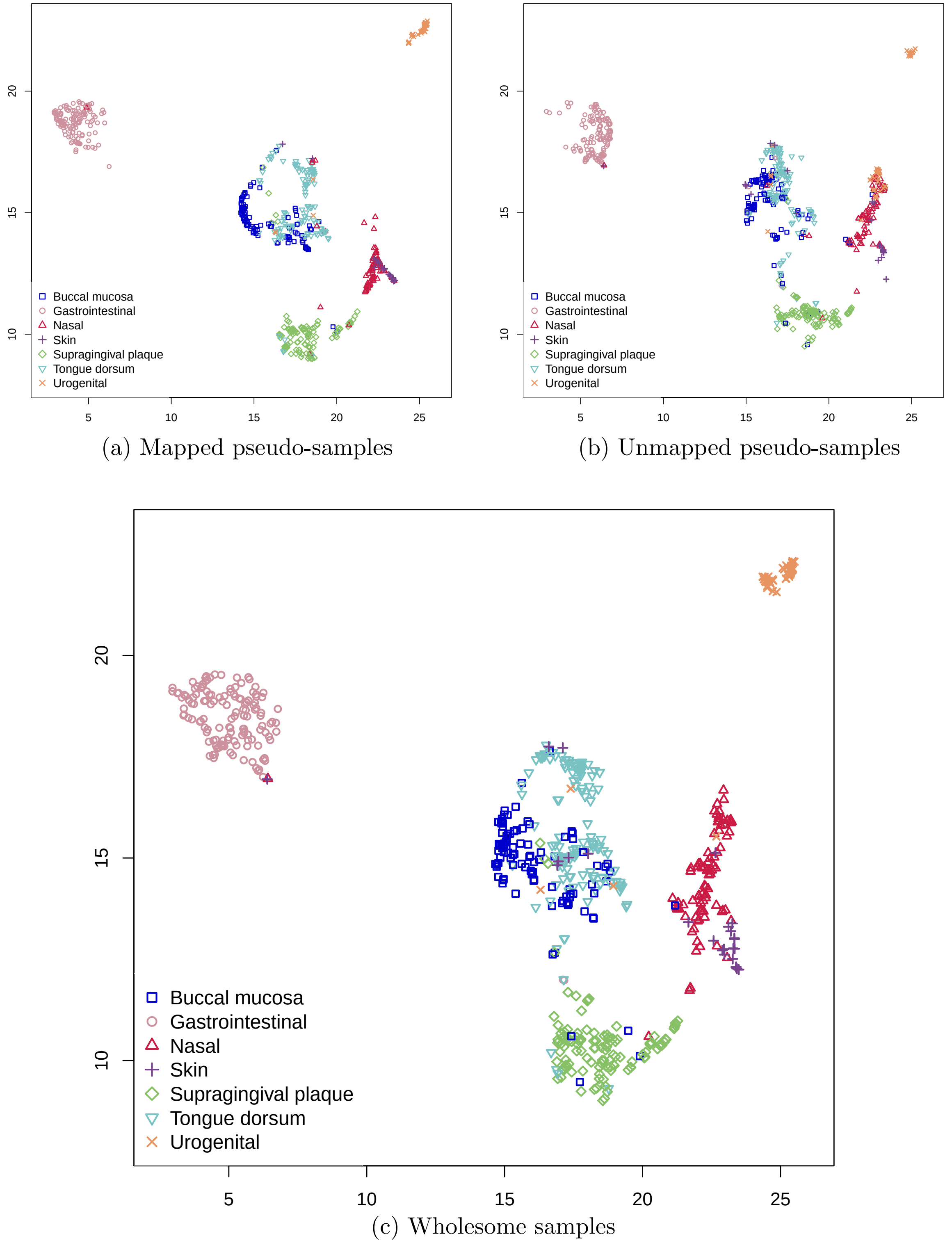


(a) Mapped pseudo-samples

(b) Unmapped pseudo-samples

(c) Wholesome samples

Figure S3: Samples clustering for mapped pseudo-samples, unmapped pseudo-samples, and wholesome samples respectively by UMAP.

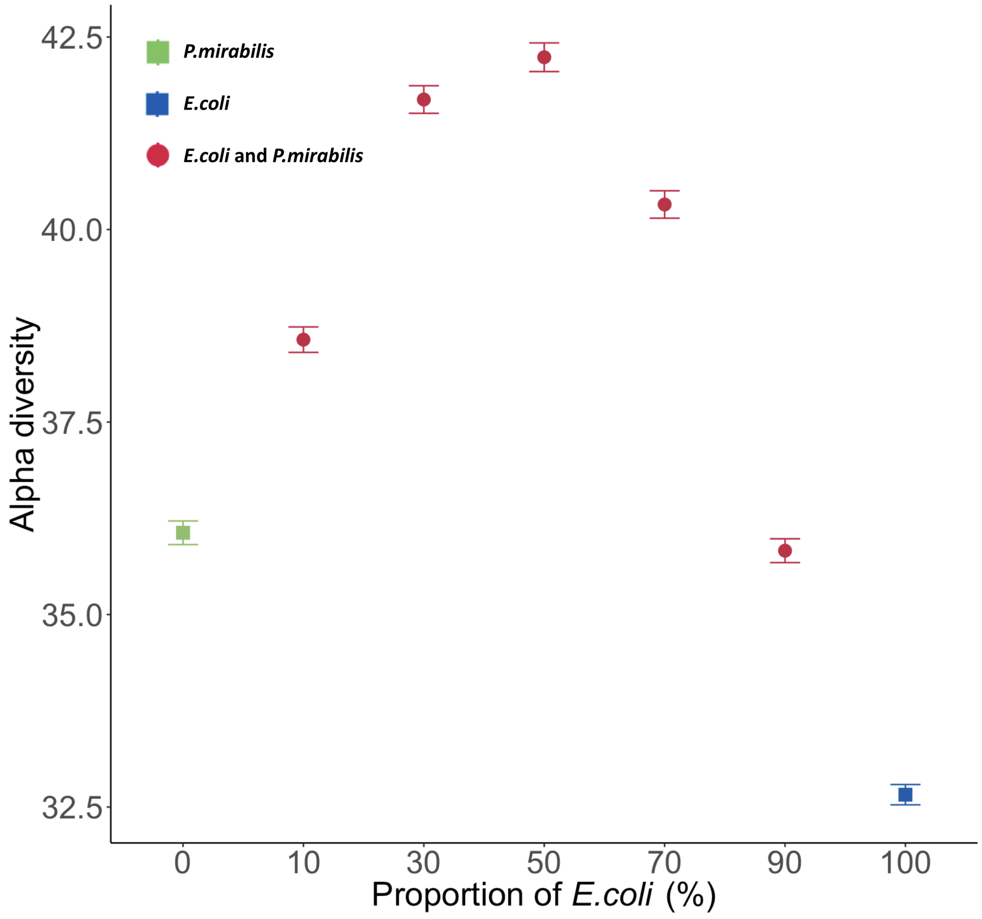


Figure S4: Within-sample diversity quantified by Nubeam using simulated samples that mixing reads at different proportions from *E. coli* and *P. mirabilis*. The bars on each dot denote ± one standard error.
